# Human transcriptional gene regulatory network compiled from 14 data resources

**DOI:** 10.1101/2021.08.28.457834

**Authors:** Vijaykumar Yogesh Muley, Rainer König

**Affiliations:** Instituto de Neurobiología, Universidad Nacional Autónoma de México, Querétaro, México; Institute for Infectious Diseases and Infection Control, Jena University Hospital, Jena, Germany; Integrated Research and Treatment Center, Center for Sepsis Control and Care, Jena University Hospital, Jena, Germany

**Keywords:** Activators, Gene expression, Gene networks, Gene regulation, Gene regulatory network, Transcription factors, Transcription regulation, Regulatory network, Repressors

## Abstract

Transcriptional regulatory network (TRN) orchestrates spatio-temporal expression of genes to generate cellular responses for survival. The transcription factors (TF) regulating expression of their target genes (TG) are the fundamental units of TRN. Several databases have been developed to catalogue human TRN based on low- and high-throughput experimental and computational studies considering their importance in understanding cellular physiology. However, literature lacks comparative assessment on the strength and weakness of each database. In this study, we compared over 2.2 million regulatory pairs between 1,379 TF and 22,518 TG assembled from 14 data resources. Our study reveals that the TF and TG were common across data resources but not their regulatory pairs. We observed that the TF and TG of the regulatory pairs showed weak expression correlation, significant gene ontology overlap, co-citations in PubMed and low numbers of TF-TG pairs representing transcriptional repression relationships. Furthermore, each TF-TG regulatory pair assigned a combined confidence score reflecting its reliability based on its presence in multiple databases and co-expression. The TRN containing 2,246,598 TF-TG pairs, of which, 44,284 with the information on TF’s activating or repressing effects on their TG is available upon request. This study brings the information about transcriptional regulation scattered over the literature and databases at one place in the form of one of the most comprehensive and complete human TRNs assembled to date, which will be a valuable resource for benchmarking TRN prediction tools, and to the scientific community working in functional genomics, gene expression and gene regulation analysis.

## Introduction

Cells have a dedicated transcriptional regulatory system, which ascertains selective expression of genes from their genomes to generate adaptive responses in a broad range of situations including response to environment, metabolic needs, cellular growth, and differentiation during development (Jacob and Monod, 1961; Laishram and Gowrishankar, 2007; Luscombe et al., 2004). The fundamental units of the transcriptional regulatory systems are transcription factors (TF) which typically bind to specific sequences in the upstream regions (promoters, enhancers) of their target genes (TG) within the genome and regulate their expression (Muley and Pathania, 2017). One TF can regulate many TG and typically several TF are needed to modulate the expression of a single gene. Many TF act synergistically on their TG and often cooperate with other TF and proteins to achieve high-fidelity transcriptional responses (Neph et al., 2012). The transcriptional regulatory system can be conceptualized in the form of a network or a graph called transcriptional regulatory network (TRN) in which nodes are TF and their TG which are connected by edges if a TF is known to regulate the TG (Babu et al., 2004; Luscombe et al., 2004). Perturbations in the TRN can lead to phenotypic diversity, initiate evolutionary adaptation and may manifest in diseases and developmental disorders (Levine and Tjian, 2003; Luscombe et al., 2004; Muley et al., 2020; Paek et al., 2016; Villar et al., 2014). Therefore, prediction and analysis of TRN has always been an active area of research in the post-genomic era.

Conventionally, transcriptional regulatory systems have been meticulously elucidated by studying TF and its interaction or binding to specific DNA sequence by biochemical and genetic experiments (Dynan and Tjian, 1983). These studies are often complemented with genome-wide expression profiling of genes in response to knockdown or knockout of a particular TF (Pathania and Muley, 2017; Salah et al., 2016). TF binding DNA sequences or motifs determined from such studies are useful in identifying TF’s novel TG by searching similar sequences or motifs in the genome, specifically in upstream regions of genes. Several databases have been set up from routinely collected potential TF binding-site motifs assembled to Position-Weight-Matrices (PWM) facilitating the prediction of new binding sites in the genome (Mathelier et al., 2014). PWM based searches allow predicting potential binding sites of TF across an entire genome (Molineris et al., 2011). However, identified binding sites can contain many false positives, which may be filtered out to some extent by confining to only binding sites found in evolutionary conserved regions (Loots and Ovcharenko, 2007).

The development of high-throughput experimental techniques such as chromatin immuno-precipitation (ChIP) followed by genome tiling array hybridization (ChIP-chip) and ChIP coupled with massive parallel sequencing (ChIP-seq) allow to experimentally obtain genomic DNA sequences bound by TF (Johnson et al., 2007; Ren et al., 2000). Large amounts of ChIP-chip and ChIP-seq data have been generated in the last decade. This data is often publicly available through several databases such as ChIP Enrichment Analysis (ChEA), hmChIP and ChIPbase (Chen et al., 2011; Lachmann et al., 2010; Yang et al., 2013). Over the years, TF-TG regulatory relationships have also been extracted from manual inspection of the literature and text mining. This information is available via databases such as TRRUST (Transcriptional Regulatory Relationships Unravelled by Sentence-based Text-mining), TFactS, the commercial database MetaCoreTM by Clarivate, and the Human Transcriptional Regulation Interactions database (HTRIdb) (Bovolenta et al., 2012; Essaghir et al., 2010; Han et al., 2015). These manually curated databases are expected to be more reliable than those derived from high-throughput experiments or predictions based on PWM analyses. However, to our knowledge, so far, no comprehensive comparative or integrative assessment of these TRN has been done. Nevertheless, Shmelkov et al. have compared the TG of seven TF across 10 commonly used pathway databases (Shmelkov et al., 2011). They observed a surprisingly small overlap between the experimentally obtained TG and TG reported in various pathway databases. Their analysis questioned the validity of transcriptional regulatory targets from many popular pathway databases, indicating need for their systematic investigation. In our survey, we found 14 human TRN available through public databases, supplementary data associated with published research articles and a commercial database called MetaCoreTM by Clarivate (Bovolenta et al., 2012; Chen et al., 2011; Essaghir et al., 2010; Gerstein et al., 2012; Han et al., 2015; Kawaji et al., 2009; Lachmann et al., 2010; Loots and Ovcharenko, 2007; Marbach et al., 2016; Mathelier et al., 2014; Molineris et al., 2011; Neph et al., 2012; Suzuki et al., 2009; Yang et al., 2013). These resources are inherently heterogeneous being obtained from a variety of experimental and computational techniques, while some are manually curated. These resources together constitute most if not all regulatory interactions between human TF and their TG, emphasizing the need for a systematic assessment to determine their reliability, and to what extent the respective database is biologically meaningful or can be used for benchmarking.

In this study, we compiled over 2,246,598 TF-TG pairs between 1,379 human TF and 22,518 TG collected from afore-mentioned 14 databases. Our analyses highlight small overlap of TF-TG pairs across databases, weak co-expression of TF and their TG. Furthermore, TF-TG pairs are well supported by co-occurrence of TF-TG in the literature, and functional overlap assessed by Gene Ontology enrichment analysis. In addition, we provide an integrative confidence score to each TF-TG pair indicative of its reliability. We have also analyzed 44,284 pairs with the information on TF’s activating or repressing effects on their TG from three databases. In summary, we provide the most up to date and complete human TRN which has wide applications in biology and is available upon request for the research community.

## Methods

### Assembly and processing of transcriptional regulatory network

The regulatory pairs between human TF and TG were obtained from the databases MetaCoreTM by Clarivate (http://thomsonreuters.com/metacore/, commercially available), TRRUST, TFactS, HTRIdb, ChEA, hmChIP, ChIPBase, ECRBase, and EdgeExpressDB (Bovolenta et al., 2012; Chen et al., 2011; Essaghir et al., 2010; Han et al., 2015; Lachmann et al., 2010; Loots and Ovcharenko, 2007; Severin et al., 2009; Yang et al., 2013). MetaCore contained several entries of TF complexes regulating a gene. Those complexes were decomposed into individual TF and each of them were paired with the listed TG. EdgeExpressDB provides TRN assembled during and from the Fantom4 project, which we could divide into two groups (Kawaji et al., 2009; Severin et al., 2009; Suzuki et al., 2009). One of which had been experimentally derived and called FantomE, which contained TF-TG pairs used as gold standard in Fantom4 study and a large number TF-TG pairs confirmed by siRNA perturbation experiments, whereas another set of TF-TG pairs was based on in-silico predictions and is referred to as FantomP (Suzuki et al., 2009). Marbach et al. predicted 32 tissue-specific TRN, which are available as supplementary data (Marbach et al., 2016). In these TRN, the TF-TG pairs contained edge weights, from which those equal or above a threshold of 0.1 were selected. These TF-TG pairs were merged into a single TRN referred to as TissueNets. Furthermore, we obtained TF-TG pairs predicted using the Total Binding Affinity (TBA) approach, which computes a TF binding profile in the gene promoter based on PWM (Molineris et al., 2011). Finally, we included a published TRN from the ENCODE project (Gerstein et al., 2012) and TRN specific to TF alone i.e. TG of the TF also encodes for TF (Neph et al., 2012).

Some of the databases contained ambiguous TF entries, which includes CACD, CST12P, ELSPBP1, GC, HACD1, HENMT1, IL10, IL6, OR5I1, PEBP1, PSMC5, TNFRSF10A, TNFRSF25, and TRIM63. These TF neither have known DNA binding domains in them nor direct evidence of DNA binding activity in the literature. Therefore, we excluded them and the remaining entries were cross-checked in databases dedicated to TF, which include Transcription Factor Classification (TFClass), DNA-binding domain (DBD), TcoF-DB databases, and the curated list of human TF reported by Vaquerizas et al. (Schaefer et al., 2011; Vaquerizas et al., 2009; Wilson et al., 2008; Wingender et al., 2013). The gene nomenclature varied across databases, which was made uniform by mapping gene synonyms to their stable NCBI Entrez identifier using the org.Hs.eg.db Bioconductor package for human genome annotation (Carlson, 2015). Finally, 1,379 TF and 22,518 TG were retained, which formed 2,246,598 unique TF-TG pairs being present in at least one of the 14 investigated databases. For a subset of TF-TG pairs, TFactS, TRRUST, and Metacore databases reported an effect of TF on its TG expression. These TF-TG pairs were represented with label “Activation” when TF found to be activated expression of its TG in literature, “Repression” if it represses expression, or both labels when TF found to activate as well as repress gene expression. The distribution of these TF-TG pairs was analyzed across 14 databases.

### Assessing the consistency of TF-TG pairs by gene ontology term overlap and co-occurrence in PubMed citation

Functional similarity between TF and TG of regulatory pairs was assessed by the overlap of their biological process Gene Ontology (GO) annotations. We collected GO terms associated with TF and TG from the org.Hs.eg.db annotation package (Carlson, 2015). The list of GO terms of the TF and the list of the GO terms of the TG were represented in a contingency table with which a Fisher’s exact test was performed testing statistical significance of the overlap. Overlaps of GO terms between TF and their TG were considered as significant if the resulting p-value was ≤ 0.05. Furthermore, we tested for co-occurrences TF and their TGs in published research articles. For this, the PubTator database was used to obtain the counts of PubMed citations in which the TF and its TG reported together (Wei et al., 2013). For each database then we computed the fraction of regulatory pairs having at least one citation mentioning both together.

### Gene expression data processing, normalization, and co-expression analysis

Transcriptomic data in the form of tags-per-million and raw counts was obtained from GTEx (phs000424.v8.p2) (GTEx Consortium, 2020). We excluded samples corresponding to Cells - EBV-transformed lymphocytes and Cells - Cultured fibroblasts to keep samples only from healthy tissues. Transcripts with raw read counts *≥* 15 in at least three samples were retained in the tags-per-million data matrix which was used for subsequent analyses. Transcripts were mapped to NCBI ENTREZ gene identifiers using org.Hs.eg.db annotation package (Carlson, 2015). Some ENTREZ identifiers were mapped to more than one transcript, from which the one with the highest average expression was retained. The dataset was quantile normalized and transformed to log2-scale by adding one as a pseudo count to expression values. This data represented the expression of 1,374 TF and their 20,370 TG in 16,704 samples, which was used to compute Pearson’s correlation coefficients (PCC) between expression profiles of TF with all genes.

### Bootstrap resampling to estimate PCC variability across 14 databases

Bootstrap resampling mimics routinely used sampling procedures in statistical analysis. Let’s assume that there is a vector *V* of absolute PCC, and *l* is the length of the *V* i.e. the number of total PCC values in *V*. When we sample *l* times PCC from *V*, each PCC value has the same chance of getting selected in each round until we have *l* number of PCC. This is called a sampling with replacement because some PCC values may get selected more than once in the random draws and the resulting sample is referred to as a resample. The collection of several resamples (generally *≥* 1,000) from the original data is called a bootstrap resampling, as if taking several random samples from the population. These lists of resamples then can be used to estimate the spread of a particular statistic compared to the observed from the original sample (Chihara and Hesterberg, 2018). In our case, we were interested in the mean statistic of PCC values of each database. Therefore, we created 10,000 resamples of the PCC of each database. The mean PCC value was computed for each resample, which forms a bootstrap distribution. The standard error of the distribution was used as the estimated variability of average PCC of the bootstrap distribution.

### Integrative confidence score assignments to transcription factors and their target genes

To compute the combined confidence score for each TF-TG pair based on the number of databases in which it was observed, we assigned weights to each database-based on the average of absolute PCC values of TF-TG pairs in them. These weights were combined to compute an integrative confidence score (iCS) for each TF-TG pair using a naïve Bayesian equation (Mering et al., 2005),

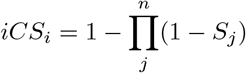

where *i* is the TF-TG pair for which the score is being calculated, *j* is the database and *n* is the number of databases reporting the *i*^*th*^ TF-TG pair. *S*_*j*_ is the weight of the *j*^*th*^ database. Since the naïve Bayesian statistic work under the assumption of independence of the databases, we used mean data weights of TissueNets, ECRbase, and TBA databases which were derived from the PWM based prediction algorithms. Likewise, data weights of Encode, hmCHIP, ChEA, and ChiPBase were also averaged since they are likely to share data from ChIP-Seq experiments. This was done to avoid redundancy in the combined score by the presence of TF-TG pairs in multiple databases just because the same data shared among them such as ChIP-seq data or based on the same technologies such as predictions based on motifs in the upstream regions of the genes.

### Evaluating functional relevance of integrative confidence scores of transcription factors and their target genes

To assess how good iCS reflect functional and citation overlap between TF and their TG, TF and its TG pair was considered as functionally linked or positive if both showed significant overlap of GO biological processes (p-value ≤ 0.05, and unrelated otherwise i.e. negative (p-value *>* 0.05). In another gold standard, a TF and its TG pair was considered as functionally linked or positive if both were mentioned together in at least one PubMed citation, and unrelated otherwise i.e. negative. For a chosen threshold iCS, the TF-TG pairs with scores greater than or equal to the threshold and belong to positive examples were classified as True Positives (TP), and those belong to negative examples are classified as False Positives (FP). Numbers of TP and FP were recorded at a series of iCS thresholds separately for GO and PubMed citation gold standards and computed positive predictive value (PPV) or precision as follows,

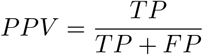

## Results and Discussion

### Concept of the study

Figure 1 shows a schematic overview of our study design and analyses. We collected known human regulatory interactions listed in 14 data resources (Figure 1A). Subsequently, we assessed the significance of overlap among the TF-TG pairs, TF, and TG across data resources. The GTEx project provides a largest human transcriptomic dataset containing 16,704 samples belonging to 52 different tissues from about 1,000 healthy individuals, which was used to compute expression PCC between TF and their TG (Figure 1B). Functional context between TF and their TG was determined using significant overlap between their Gene Ontology terms, and indirectly based on their co-occurrence in PubMed citations (Figure 1C). Each TF-TG pair was assigned an integrative confidence score considering its presence in multiple data resources and quality of them (Figure 1D).

**Figure 1:**
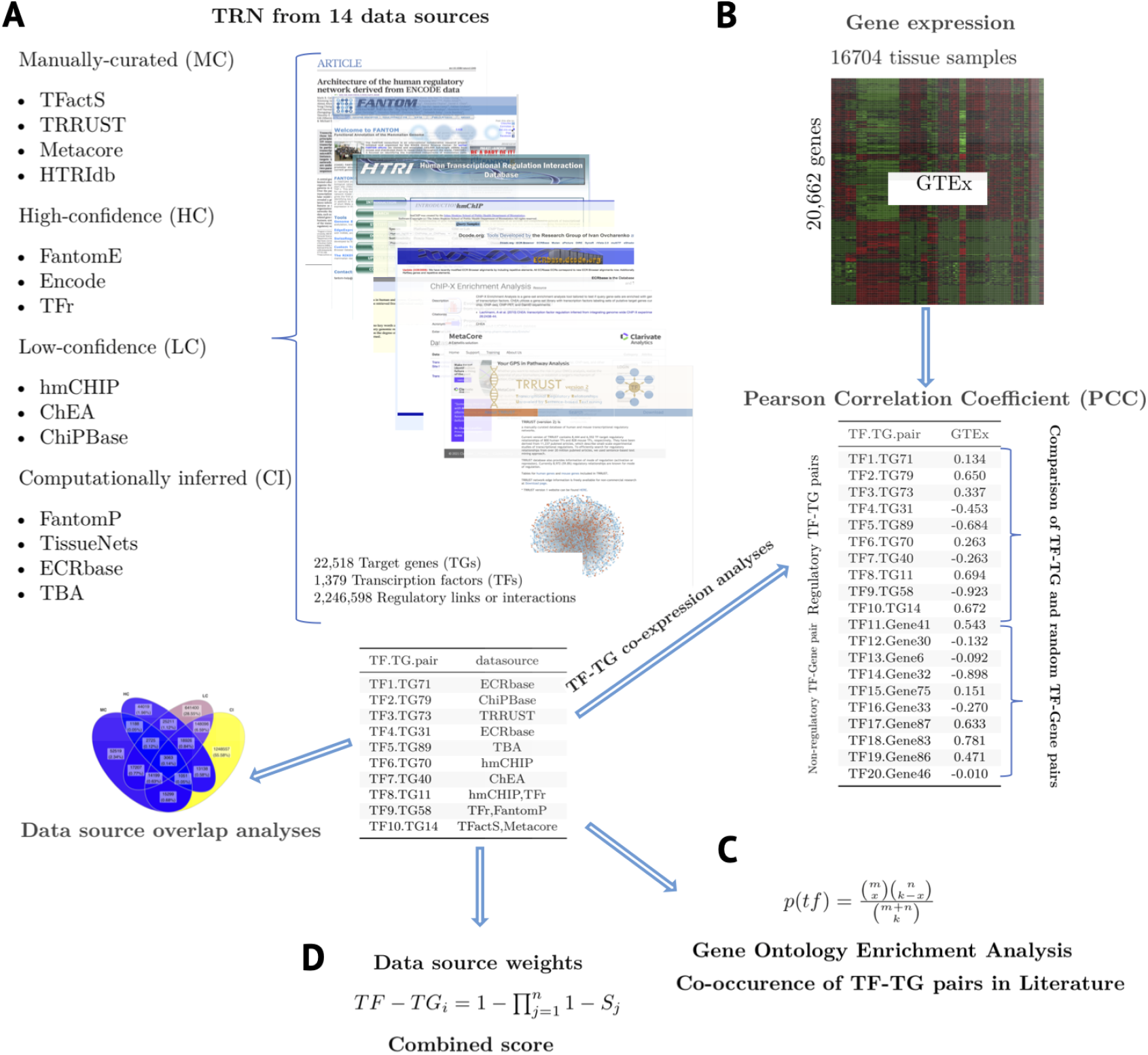
The workflow. A) Transcriptional regulatory networks from 14 databases were assembled and categorized into four groups according to their quality and data type. B) Co-expression analysis of transcription factors (TF) and their target genes (TG) using 16,704 transcriptomes obtained from GTEx web-portal. C) Analysis of Gene Ontology term overlap and co-citations in PubMed between TF and their TG. D) Assignement of a combined confidence score to each TF-TG pairs reflecting its quality and reliability.

### A quality-based categorization of the regulatory interactions

We compiled 2,246,598 TF-TG pairs of 1,379 TF and 22,518 TG from 14 databases (Table 1. TF were cross-checked from three TF repositories and a curated list of human TF from a published study to exclude ambiguous entries (Supplementary figure 1) (Schaefer et al., 2011; Vaquerizas et al., 2009; Wilson et al., 2008; Wingender et al., 2013). The quality and the number of TF-TG pairs varied across 14 databases due to the differences in experimental and computational methods used to derive them. These databases were grouped into four categories for brevity (Table 1). The TRRUST, TFactS, HTRIdb, and MetaCore databases comprise manually inspected 107,251 TF-TG pairs confirmed in low-throughput experimental studies and were obtained from literature review, which were referred together as the Manually Curated (MC) (Bovolenta et al., 2012; Essaghir et al., 2010; Han et al., 2015). The second category contained 109,321 pairs obtained from high-throughput experimental studies conducted as part of the projects Fantom4 and ENCODE, and also TF-TG pairs of 531 TF which were identified in 41 cells and tissues types using in vivo DNaseI footprints, hereafter referred to as FantomE, Encode, and TFr, respectively (Gerstein et al., 2012; Neph et al., 2012; Severin et al., 2009). These databases were collectively referred to as High-Confidence (HC) since they were rigorously analyzed in the associated studies to remove spurious TF-TG pairs. For example, FantomE represents TF-TG pairs which were derived from ChIP-chip experiments and some of them were further confirmed by siRNA knockdown studies in the human monocyte cell line THP-1 (Suzuki et al., 2009), TF-TG pairs from ENCODE project were assembled from ChIP-seq experiments of 125 TF in five different cell lines (Gerstein et al., 2012). The 870,827 TF-TG pairs were obtained from ChIPbase, ChEA, and hmChIP databases, which were referred to as Low confidence (LC), since most data was derived from published ChIP-chip and ChIP-seq studies without rigorous follow up screening (Chen et al., 2011; Lachmann et al., 2010; Yang et al., 2013). The fourth, Computationally Inferred (CI) category contained 1,462,329 TF-TG pairs from four databases referred to as FantomP, ECRbase, TBA, and TissueNets. These TF-TG pairs were inferred by searching PWM of known TF in the regulatory regions of the TG. FantomP included TF-TG pairs computationally predicted as a part of the Fantom4 project and based on the conservation of TF binding sites (TFBS) within regions identified by deepCAGE which bases on explaining the expression of the gene employing motif activity analysis of its promoter (Suzuki et al., 2009). The Transcription factor Binding Affinity (TBA) method quantifies the binding affinity of a TF for the whole promoter of its TG using a sliding window (Molineris et al., 2011). ECRbase TF-TG pairs were identified based on the presence of TF binding sites in evolutionarily conserved promoter elements of vertebrates (Loots and Ovcharenko, 2007). TissueNets contains TF-TG pairs from tissue specific TRN inferred by integrating binding site motifs of TF with promoter and enhancer activity data from the Fantom5 project (Marbach et al., 2016). These CI databases comprised 65% of all investigated TF-TG pairs.

**Table 1:** A description of the 14 transcriptional regulatory networks analyzed in this study

| Database | # of TF-TG pairs |  | # of TF | # of TG | Average node degree |  | Database/Reference |
| --- | --- | --- | --- | --- | --- | --- | --- |
|  | Original* | Processed† |  |  | TF (in) | TG (out) |  |
| Manually curated (MC) |  |  |  |  |  |  |  |
| TFactS | 4254 | 4206 | 272 | 1880 | 15.46 | 2.24 | TFactS |
| TRRUST | 8013 | 7984 | 746 | 2358 | 10.70 | 3.39 | TRRUST |
| Metacore | 57033 | 56294 | 976 | 10631 | 57.68 | 5.30 | Metacore |
| HTRIdb | 48993 | 48765 | 277 | 16662 | 176.05 | 2.93 | HTRIdb |
| High-Confidence (HC) |  |  |  |  |  |  |  |
| FantomE | 40549 | 40318 | 168 | 13315 | 239.99 | 3.03 | EdgeExpressDB |
| Encode | 24629 | 24532 | 116 | 8230 | 211.48 | 2.98 | Gerstein et al., 2012 |
| TFr | 47222 | 47085 | 531 | 503 | 88.67 | 93.61 | Neph et al., 2012 |
| Low-Confidence (LC) |  |  |  |  |  |  |  |
| hmCHIP | 355762 | 355409 | 69 | 14257 | 5150.86 | 24.93 | hmCHIP |
| ChEA | 344056 | 339901 | 198 | 19546 | 1716.67 | 17.39 | ChEA |
| ChIPBase | 415441 | 414886 | 79 | 18206 | 5251.72 | 22.79 | ChIPBase |
| Computationally inferred (CI) |  |  |  |  |  |  |  |
| FantomP | 229745 | 229205 | 305 | 9540 | 751.49 | 24.03 | EdgeExpressDB |
| TissueNets | ND | 177439 | 583 | 14252 | 304.36 | 12.45 | Marbach et al., 2016 |
| ECRbase | 1081089 | 1001955 | 174 | 14956 | 5758.36 | 66.99 | ECRbase |
| TBA | 187765 | 187338 | 125 | 20772 | 1498.70 | 9.02 | Molineris et al., 2011 |

*Note:*

TRN is considered as a directed network in which in-degree represents the number of TF regulating a particular gene, and out-degree is the number of TG a particular TF regulates. Interestingly, average in-degree and out-degree is substantially lower in MC and HC databases comapred to LC and CI (Table 1). Figure 2A shows that only 3,063 (0.14%) TF-TG pairs were common in all four categories. The CI and LC databases represent 55.58% and 28.55% unique TF-TG pairs, respectively. The statistical significance of overlap between these four categories was analyzed using a Fisher’s exact test (Figure 2B). The CI category does not show a statistically significant overlap of TF-TG pairs with any other category. Overall, these results show that common TF-TG pairs were rather underrepresented across categories, even though TG were common in them, and to some extent, TF (Figure 2B).

**Figure 2:**
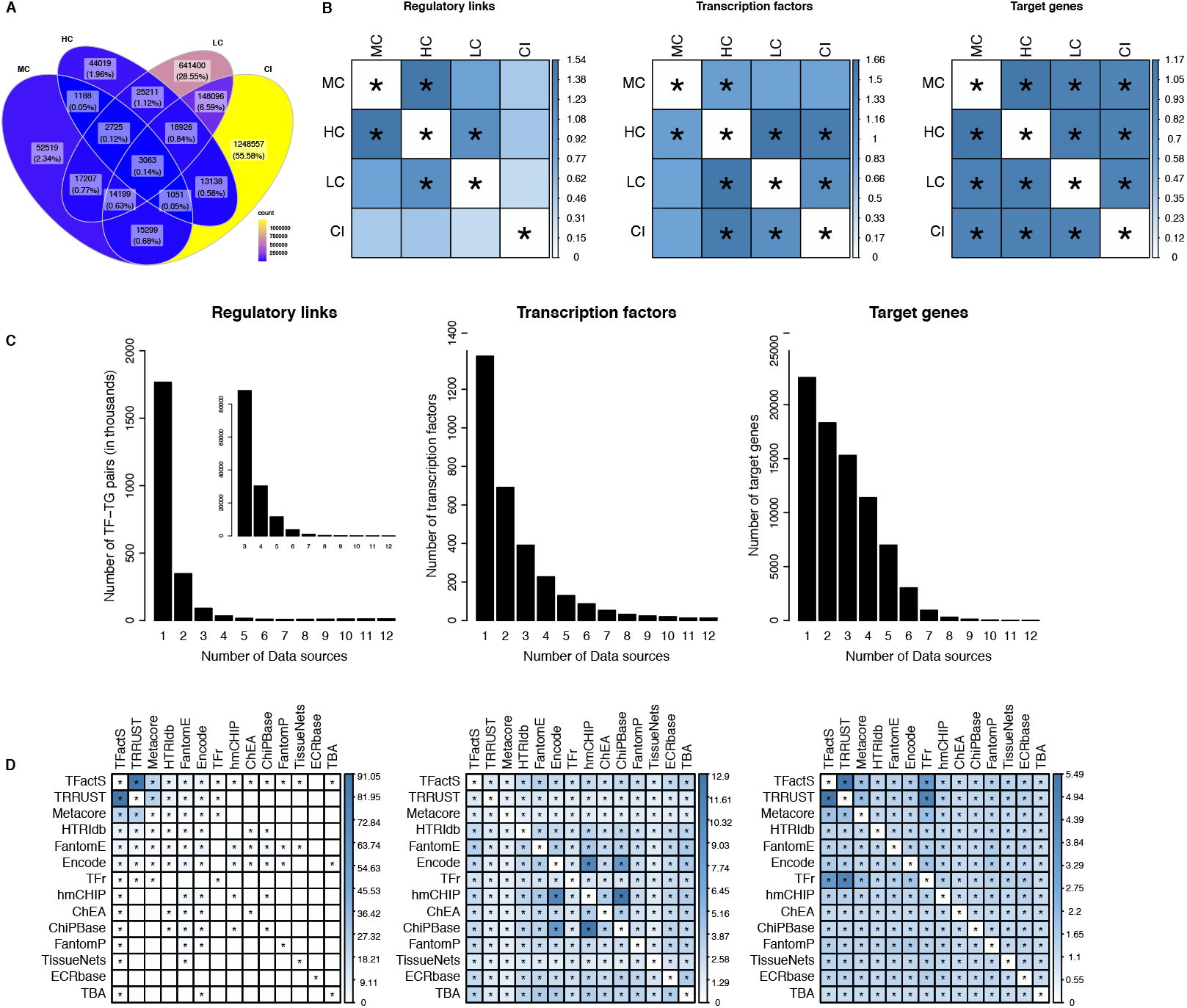
Regulatory interactions are very differently distributed, but transcription factors (TF) and their target genes (TG) show significant overlap across 14 databases categorized into four groups. A) The venn diagram shows the distribution of 2,246,598 reported TF-TG pairs across the four categories as manually curated (MC), high confidence (HC), low confidence (LC), and computationally inferred (CI). B) Heatmaps show the statistical significance of the overlap of TF-TG pairs, TF and TG. The colour of the boxes codes for the significance of overlap using odds ratios. Asterisk mark indicates a significant overlap between the categories (Fisher’s exact test p-value ≤ 0.05). C) Co-occurrence frequencies of TF-TG pairs, TF, and TG across 14 databases. The inset displays the frequency distribution of reported TF-TG pairs on a smaller scale, illustrating that commonly recorded TF-TG pairs but rarely covered in more than 6 databases. D) Legend reads the same as in panel B for the 14 databases.

### Regulatory interactions are very differently distributed, but transcription factors and target genes show significant overlap across 14 databases

The coverage of TF varied across 14 databases ranging from 69 in the hmChIP to 976 in Metacore, while TG coverage was highest in the TBA dataset (20,772 genes) and was lowest in TFr with 503 TG, which all encode TF (Table 1). The average TG was 11,793 across all databases. The combined TF-TG pairs from 14 databases contained 1,379 unique TF, which substantially exceeded the highest number of TF in the Metacore database. Furthermore, this figure is close to 1,391 high-confidence sequence-specific TF-coding genes identified and verified by manual inspection in humans (Vaquerizas et al., 2009). Altogether, these numbers indicate that the coverage of TRN assembled from 14 databases is much better than their individual sources.

The number of TF-TG pairs varied highly across databases. ECRbase alone contributed to about 45% of all assembled pairs. Figure 2C shows that the same TF-TG pairs are rarely present in more than two databases but TG in several, and to some extent TF as well. Surprisingly, 79% of all TF-TG pairs were unique to individual databases, while 15% observed in two out of 14, and only 6% TF-TG pairs were common to more than three databases (Figure 2C and Inset). Figure 2D shows a statistically significant overlap among TF or TG observed across all databases, while mostly insignificant for TF-TG pairs across 14 databases except for MC, as assessed by Fisher’s exact test. Notably, LC and CI databases did not show a significant overlap with MC databases. In contrast, MC databases showed significant overlaps among them, and with HC (Figure 2D). Few databases showed significant overlap with several other databases, which are - TFactS (with 12 databases out of 13), FantomE (with 10 out of 13), and Encode (with 10 out of 13), suggesting the high-quality TF-TG pairs in them. Overall, TF and TG were significantly common across various databases but not the TF-TG pairs between them.

### Functional overlap and co-citation of transcription factors and their target genes

TF orchestrate controlled expression of genes in response to intra- and extra-cellular signals. Genes expressed in response to a particular signal are often functionally related (Muley and Ranjan, 2013). Therefore, TF and their TG, being the part of the same signaling response, are expected to have functional overlap. Fisher’s exact test was used to evaluate independence of GO biological process of the TF and the TG constituting each regulatory pair (Ashburner et al., 2000). Figure 3A shows percentages of TF and TG pairs with significant p-values (≤ 0.05) in each database. MC except HTRIdb and HC databases showed higher fractions of TF-TG pairs with significant overlap compared to LC and CI databases. The best percent coverage was observed for the TFr database. Presumably, a higher GO term overlap was borne from the nature of TG, which all encode for TF. The TFr network was reconstructed to understand how one TF regulates expression of another TF (Neph et al., 2012). Since both TF and TG in this network are TF, they are likely to share overlapping GO terms related to transcriptional regulation.

**Figure 3:**
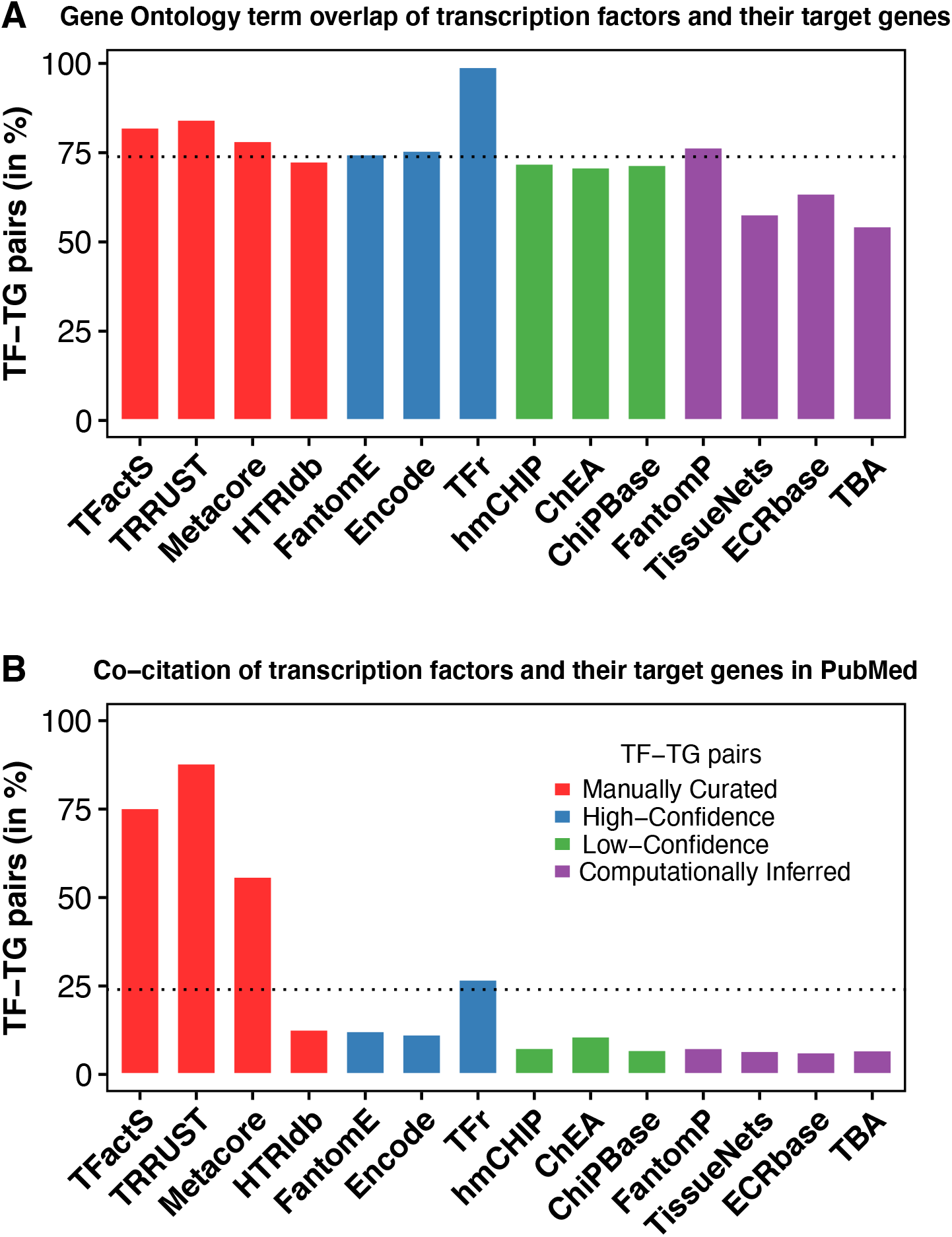
An overlap of biological process Gene Ontology (GO) and PubMed citations of transcription factors (TF) and their target genes (TG) across 14 databases. Figure shows the percentage of total TF-TG pairs with significant overlap of GO biological processes between their TF and TG (A) and the presence in the same PubMed citation (B). Manually curated databases show higher GO term overlap and PubMed co-citations indicating the reliability of TF-TG pairs therein.

The guilt-by-association approach was also used to check whether TF and its TG are related. The PubTator database provides access to genes mentioned in millions of citations available at PubMed (Wei et al. 2013). In each database, we computed a fraction of TF and TG of regulatory pairs observed both at least in one PubMed citation. Only four out of 14 databases had coverage of TF-TG pairs overlapping citation for more than 25% of their total TF-TG pairs (Figure 3B). Of them, TFactS, TRRUST, and Metacore databases contain more than 50% TF-TG pairs that are found together at least in one citation, which was expected since these databases contain manually curated TF-TG pairs from literature. However, HTRIdb database claims to contain experimentally verified transcriptional regulatory interactions (Bovolenta et al., 2012), but our analysis shows more than 85% of its TF-TG pairs do not appear in the same citations in PubMed. Its coverage was similar to LC and CI databases, which indicates less reliable regulatory pairs in HTRIdb (Figure 3B). It also has low GO overlap between TF-TG pairs compared to remaining databases in MC category (Figure 3A). The TFr is the fourth database which had about 25% of its TF-TG pairs in the same citations. Presumably, TF and TG of this database co-occurring in studies related to TF since TG encodes for TF in this network.

Overall, databases with computationally inferred TF-TG pairs showed poor GO overlap compared to other databases, and PubMed citation overlap could be observed only for manually curated databases.

### Weak expression correlation between transcription factors and their target genes

Co-expression is a good indicator of regulatory relationship since TF often correlated at expression level with their TG than with other random genes (called hereafter TF-Gene pairs) (Muley and Ranjan, 2013). This prompted us to explore to what extent TF and their TG related at expression level, whether it differs with the TF-Gene pairs. The Pearson’s correlation coefficients (PCC) were computed between TF and all genes using GTEx data of 16,704 transcriptomes of 52 tissues. Computed PCC values showed normal distribution (Supplementary figure 2). The PCC values were transformed to their absolute values since negative PCC values can be borne from inhibitory effects of TF on their TG, while positive when they activate their expression. One-sided Wilcoxon-test indicated that the absolute PCC values of TF-TG pairs were statistically significantly higher than the random TF-Gene pairs (Figure 4A).

**Figure 4:**
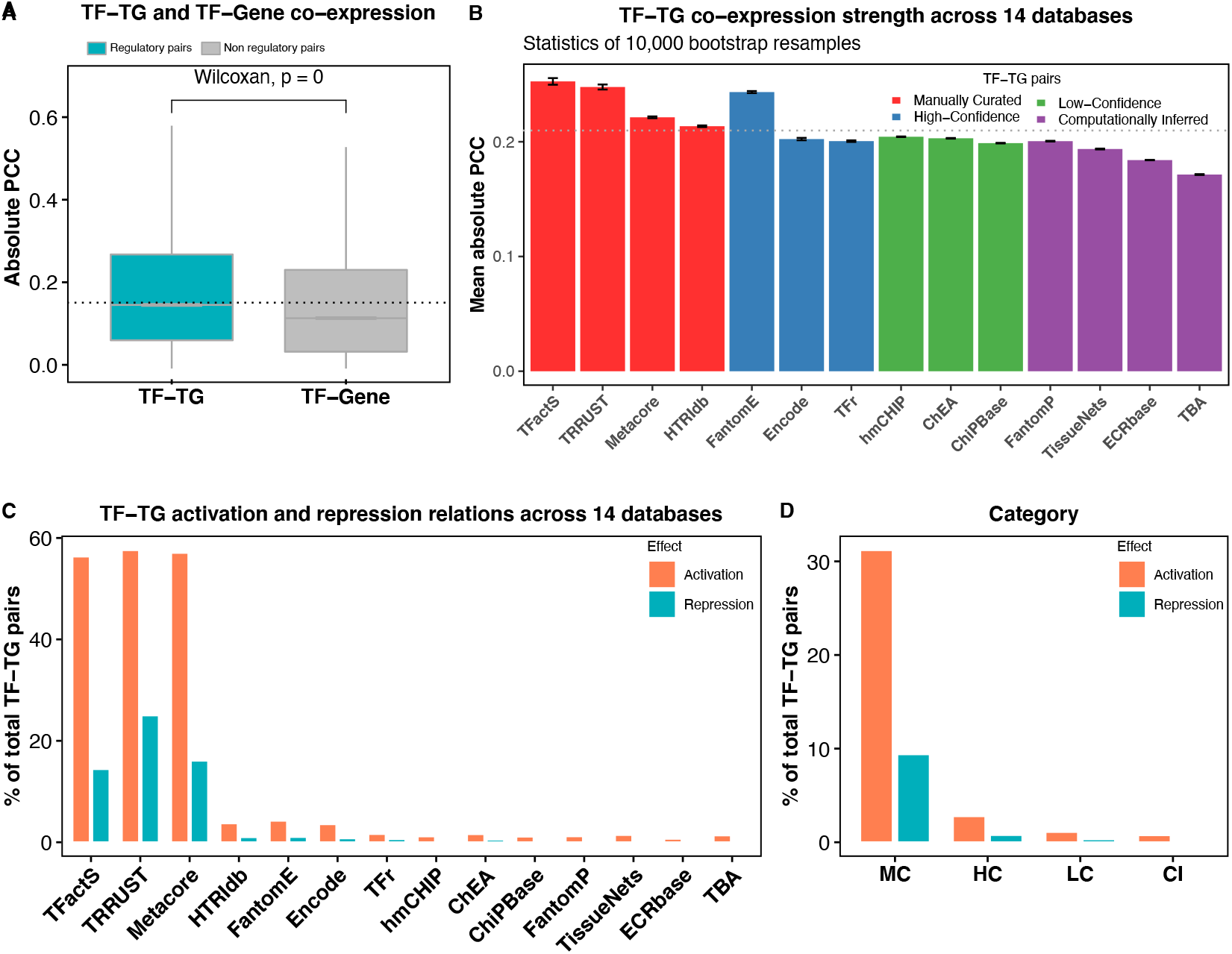
Co-expression analysis of transcription factors (TF) and their target genes (TG) reveals weak expression correlation between them. Panel A) displays the distributions of absolute values of Pearson’s correlation coefficients (PCC) between TF and their TG from TF-TG pairs (green) and non-TF-TG pairs of TF and genes (grey). B) Bootstrap distributions of average PCC values from 10,000 resamples. C) The percentage of total TF-TG pairs with TF having activating (Activation) or repressing (Repression) effect on their TG across 14 databases, and across four categories (D) i.e. manually curated (MC), high confidence (HC), low confidence (LC), and computationally inferred (CI).

However, the median PCC score was around 0.2 suggesting a weak expression correlation between TF and their TG.

To get better idea on PCC values assigned to TF-TG pairs in each database, we used simulation based bootstrap resampling procedure (Chihara and Hesterberg, 2018). A bootstrap distribution of mean PCC values was created from 10,000 resamples created from by randomly drawing original PCC values with replacement that were associated with TF-TG pairs of each database (Supplementary figure 3). The idea was that the original sample approximates the population from which it was drawn. So, resamples from the original PCC scores approximate what we would observe if we drew many random samples from the population. Figure 4B shows that the mean PCC and standard error of bootstrap distributions vary across 14 databases (please refer to Supplementary figure 4 for distribution plots). The average PCC estimates were substantially higher for MC databases and lower as the database quality decreased with a couple of exceptions (Figure 4B). For instance, HTRIdb has a mean PCC lower than some HC and CI databases. In turn, the mean PCC of FantomE was as good as the MC databases. Notably, the lowest and the second lowest scores were for the TBA and ECRbase databases, respectively (Figure 4B). This is interesting because TF-TG pairs in TBA were predicted by assessing only binding site motifs of TF in the promoter regions of the genes. In contrast, the ECRbase also uses this kind of information but the resulting TF binding motifs were confirmed by investigating other mammalian genomes, indicating reliability of its TF-TG pairs than TBA, which is also reflected in higher mean PCC value than TBA. These results suggest that the higher PCC scores of TF-TG pairs correlate with the quality of databases in which they exhibit and are more likely to represent true regulatory interactions. These results further confirm the higher expression correlation between TF-TG pairs compared to the non-regulatory TF-gene pairs.

### Regulatory pairs with repressive effect of transcription factors on their target genes are underrepresented compared to pairs with activating effect across 14 databases

TFactS, TRRUST, and Metacore databases provide information on the effect of TF on it’s TG expression manually inspected in the literature. We compiled this information and found 44,284 TF-TG pairs in which 33,873 pairs were composed of TF activating its TG, 9,821 where TF repressing its TG, and 590 pairs where TF was shown to activate as well as repress its TG. The coverage of these TF-TG pairs was plotted across databases and their four categories. Obviously, the fraction of TF-TG pairs was high in TFactS, TRRUST, and Metacore databases. Interestingly, we observed substantially low numbers of TF-TG pairs in which TF represses its TG, while TF-TG pairs with activating effect were abundant across all databases (Figure 4C). This effect was more pronounced when plotted for four categories (Figure 4D). Transcriptional repression in general is more complicated than activation to investigate at experimental level (Johnson, 1995). Our results suggest that the TF-TG pairs exhibiting repression relationships are less explored experimentally or underrepresented in the literature.

### Assigning combined scores to TF-TG pairs by accounting for presence across 14 databases and their quality

The heterogeneous quality of TF-TG pairs across 14 databases pose the challenge in trusting whether a listed TF-TG regulatory interaction takes place in the cell. To identify trustworthy TF-TG pairs, the mean PCC score assigned to TF-TG pairs of each database was used as its quality weight to compute an integrative confidence score (iCS) employing a naïve Bayesian approach (details in Methods). The iCS score ranges from 0 to 1 and higher scores of TF-TG pairs suggest their presence in multiple databases with higher weights. The number of TF-TG pairs drastically decrease over 0.2 threshold of iCS (Figure 5A). It suggests that the TF-TG pairs with iCS cutoff 0.2 or above are more reliable and evident in multiple databases. To quantify this systematically, we computed performance measures which have been routinely used to validate protein-protein interaction methods (Muley and Ranjan, 2012). Here, we assumed that the TF-TG pair is reliable if a TF and its TG are functionally linked. Then,we should observe more functionally linked TF-TG pairs at higher iCS thresholds than not functionally linked. Therefore, the functional linkage of TF-TG pair was considered if TF and TG showed significant GO overlap or found together at least in one PubMed citation and treated them as positive examples, otherwise negatives. Then true positives (TP) and false positives (FP) were recorded at a series of iCS thresholds. The TF-TG pairs belonging to positive examples with their iCS values equal to or above a particular iCS threshold were treated as TP, and those belonging to negative examples as FP. Figure 5B shows precision (positive predictive value) observed at a series of iCS thresholds for GO and PubMed citation overlap separately. As expected, precision increases with the increase in iCS threshold suggesting that higher number of TF-TG pairs with GO or PubMed citation overlap. These results indicate that the iCS score can be used as an indicator not only of presence of TF-TG in multiple databases but also their functional linkage in the assembled TRN.

**Figure 5:**
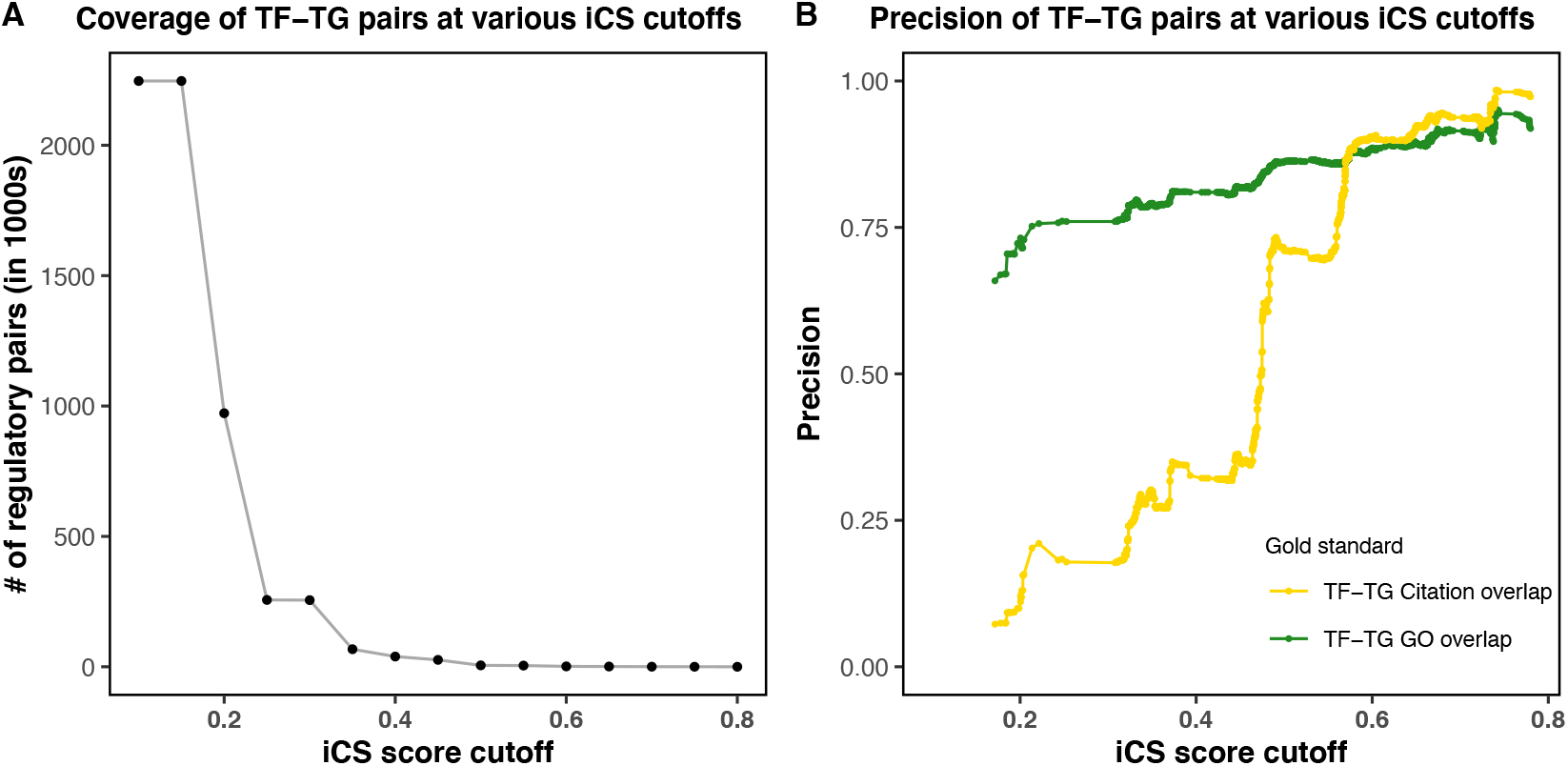
Integrative confidence scores (iCS) assigned to transcription factors (TF) and their target genes (TG) reflect the reliability of regulatory interactions at functional and co-citation levels. Each point on Panel (A) represents the number of TF-TG pairs above a series of iCS threshold given at X-axis. Panel (B) depicts positive predictive value or precision using GO and PubMed citation gold standards recorded at a series of iCS thresholds given at X-axis. Pnael shows at higher precision increases at higher iCS cutoffs suggesting higher numebr of TF-TG pairs with GO or PubMed citation overlap i.e. functionally linked.

## Conclusions

In this study, we compiled the largest human TRN available to date from 14 databases. Based on analyses presentated in this study, it can be concluded that the regulatory pairs are inconsistent across databases and TF-TG consituting them show weak expression correlations, high functional overlap TF and rarely cited together in the same PubMed citation. TfactS, TRRUST and Metacore databases were exception to this trend and represented high quality regulatory pairs considering high functional overlpap, co-expression, and co-citation between TF and their TG in them. We also provide integrative confidence scores to each TF-TG pair reflecting its reliability. Presumably, this combined TRN will be useful in benchmarking of prediction tools (Marbach et al., 2012; Muley, 2021) and help the research community in understanding transcription regulation at local and global level.

## Data availability

The data compiled in this study is available upon reasonable request to the authors for now, and will be made available freely after publication of this study in peer-reviewed journal.

## Acknowledgements

Authors acknowledge Alexandra Poos, Ph.D. for her kind help in providing two datasets used in this work, and Luis Alberto Aguilar Bautista, Laboratorio Nacional de Visualización Científica Avanzada (LAVIS) for technical support. The Genotype-Tissue Expression (GTEx) Project was supported by the Common Fund of the Office of the Director of the National Institutes of Health, and by NCI, NHGRI, NHLBI, NIDA, NIMH, and NINDS. The data (GTEx analysis V8) used for the analyses described in this manuscript were obtained from the GTEx Portal on 01/30/2020.

---

**Supplementary figure 1:**
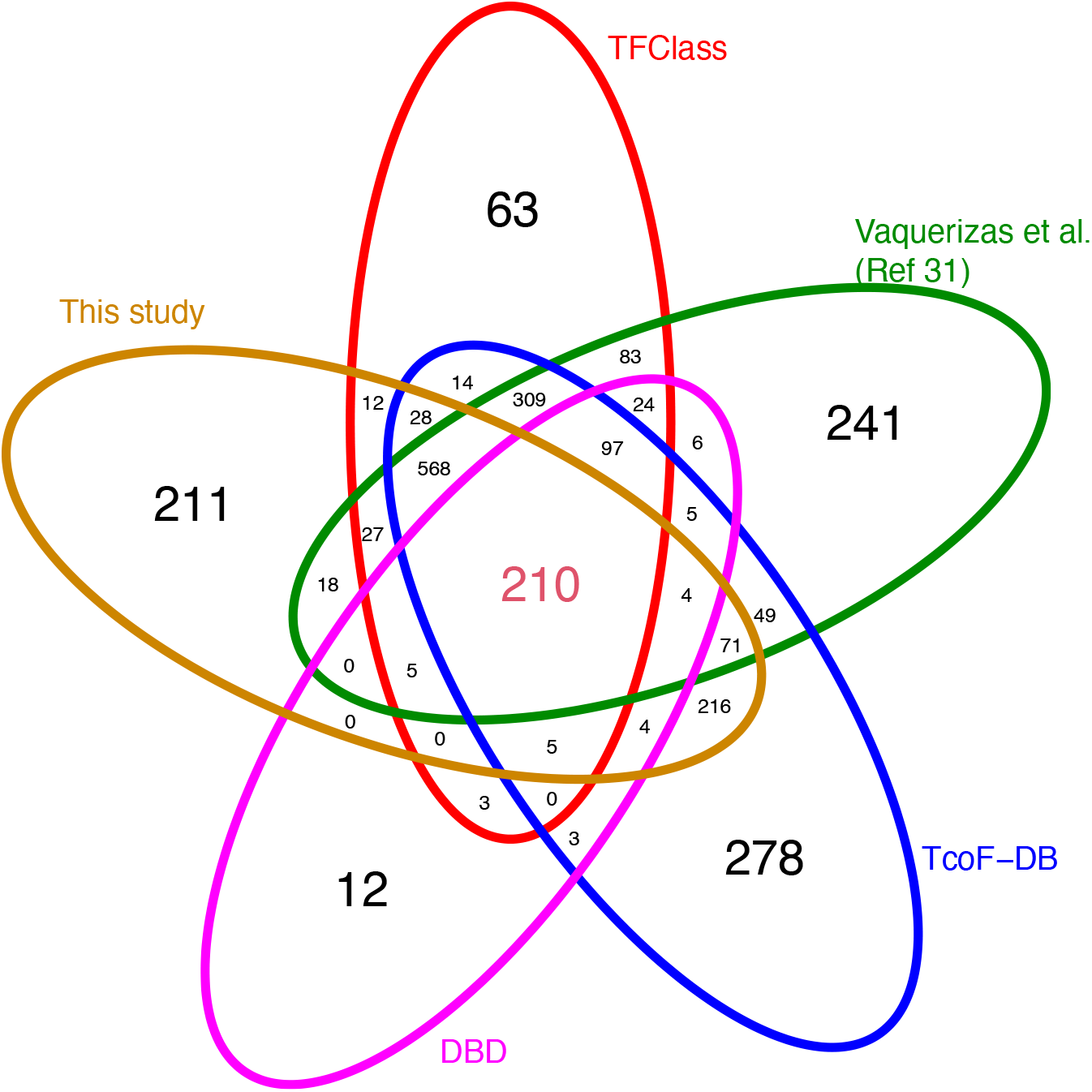
The distribution and overlap of transcription factors obtained from five resources. The venn diagram shows the distribution of transcription factors (TF) from four resources and its comparison with TF selected from regulatory pairs assembled in this study.

**Supplementary figure 2:**
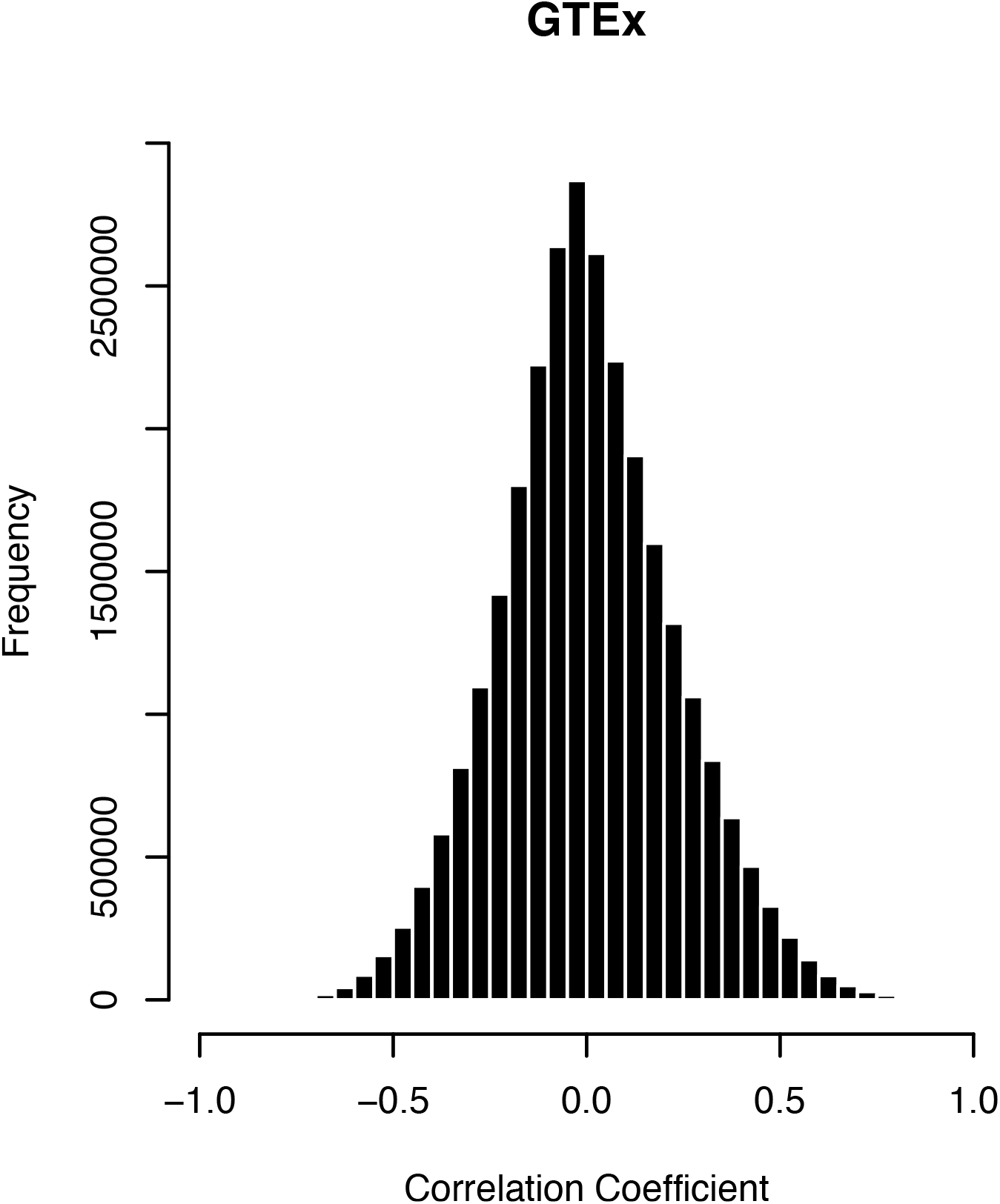
The distribution of Pearson’s expression correlation coefficients between transcription factors and analyzed genes using GTEx transcriptomic dataset.

**Supplementary figure 3:**
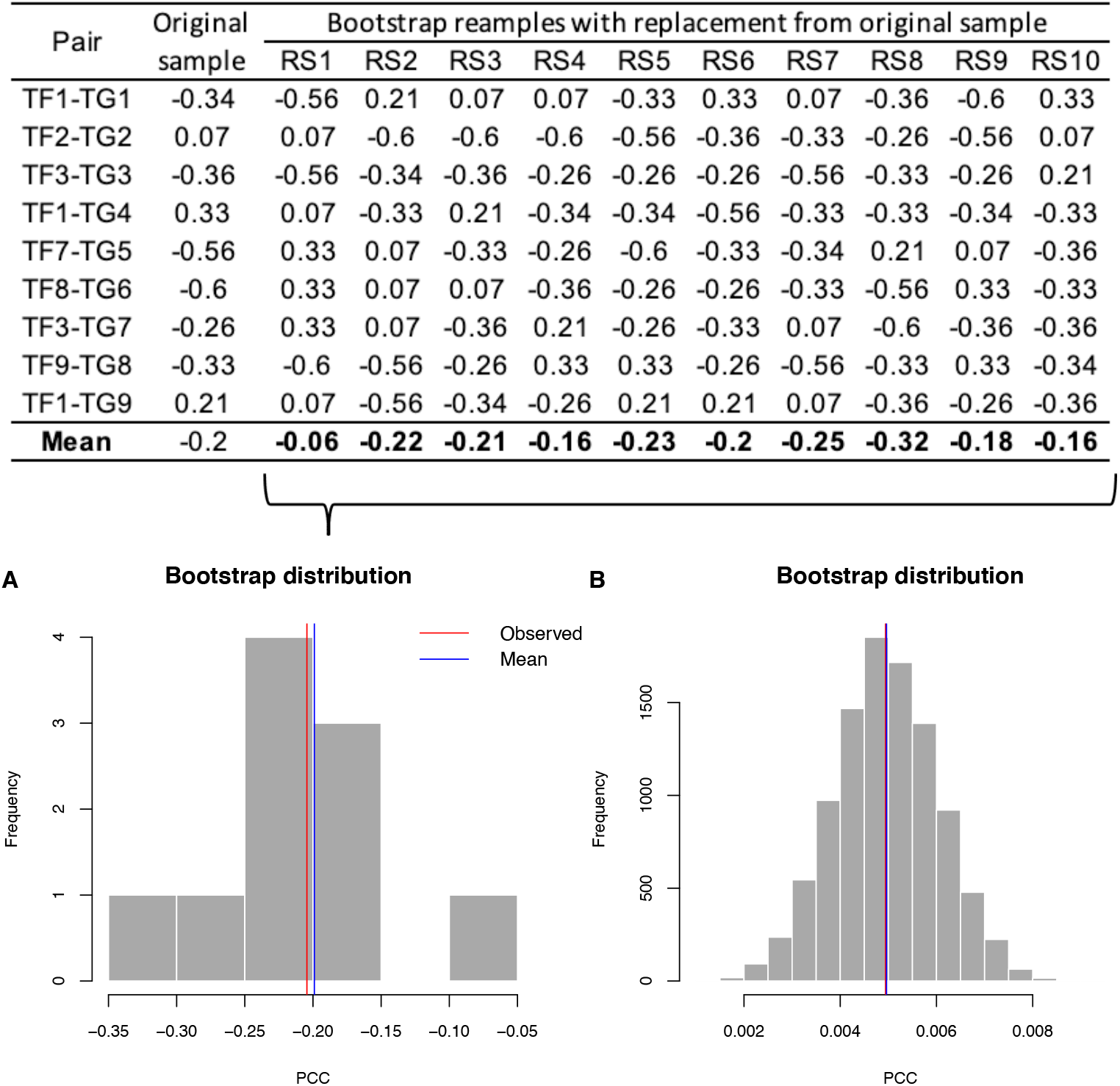
A demonstration of bootstrap resampling procedure using a toy example. A toy example showing transcription factors (TF) and their target genes (TG) in the first column, while the second column (original sample) represents expression similarity between TF-TG measured as Pearson correlation coefficient (PCC). These observed PCC were randomly drawn with replacement to create 10 bootstrap resamples (Columns RS1-RS10), and computed their means, which form a bootstrap distribution as shown in (A). The mean of the bootstrap distribution and the observed PCC (mean of the original samples) is shown by vertical bars. Panel B shows an ideal bootstrap distribution of means of 10,000 bootstrap resamples created using 25,000 expression correlations between non-regulatory gene pairs. It shows a normal distribution and the observed mean and mean of the bootstrap distributions is close. The spread of the distribution indicates the variability of PCC in the original data. In all our analyses, absolute PCC were used to create 10,000 bootstrap resamples and reported their means and standard errors.

**Supplementary figure 4:**
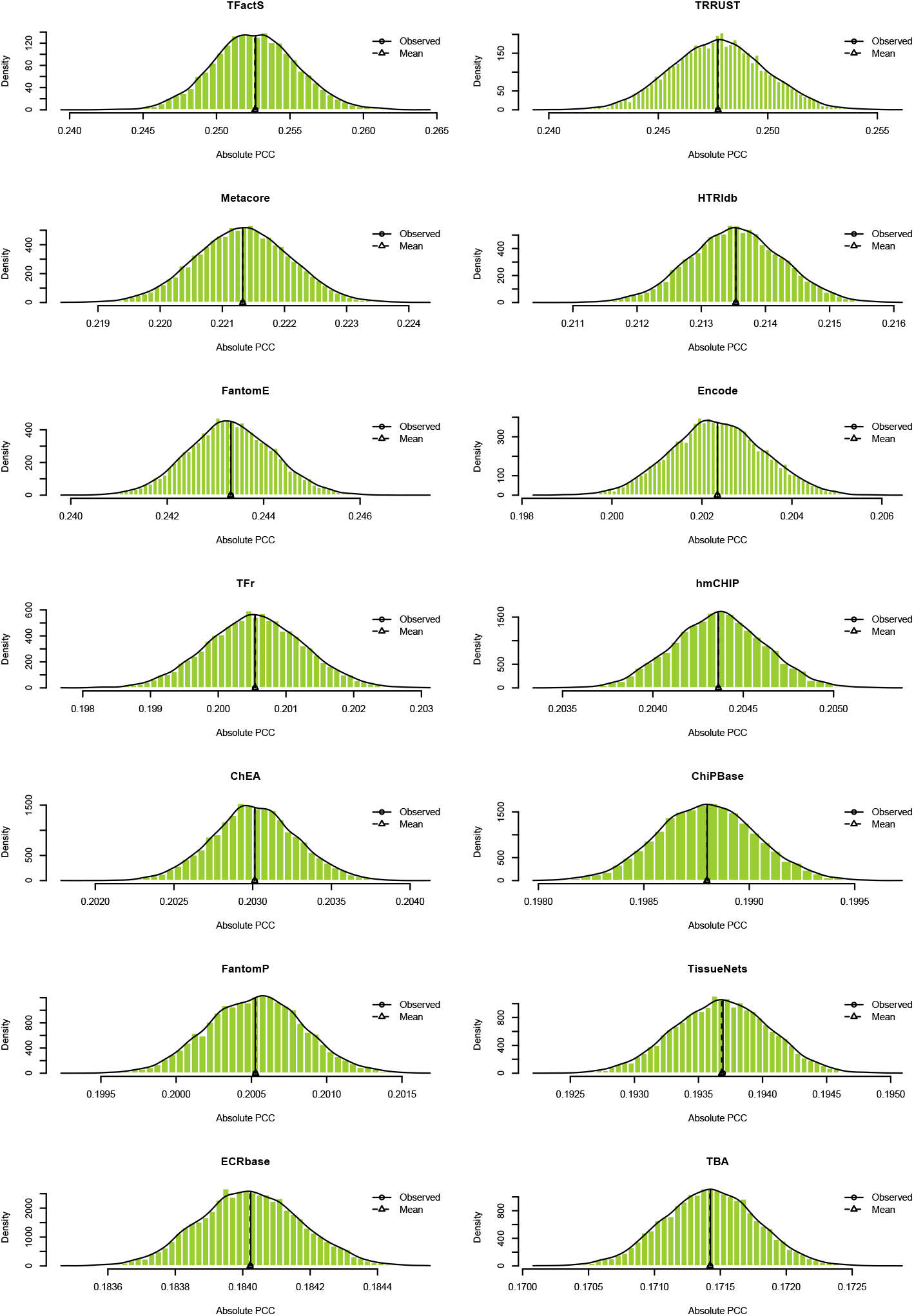
Bootstrap distributions created from 10,000 resamples of PCC associated with transcription factors and their target genes from each of the 14 databases. The observed mean and the bootstrap distribution mean shows overlap suggesting that the observed mean approximates the sampling mean.

